# Bdnf-Ntrk2 Signaling Promotes but is not Essential for Spinal Cord Myelination in Larval Zebrafish

**DOI:** 10.1101/2025.02.19.639062

**Authors:** Kristen Russell, Christina A. Kearns, Macie B. Walker, Christopher S. Knoeckel, Angeles B. Ribera, Caleb A. Doll, Bruce Appel

## Abstract

Myelin, a specialized membrane produced by oligodendroglial cells in the central nervous system, wraps axons to enhance conduction velocity and maintain axon health. Not all axons are myelinated, and not all myelinated axons are uniformly wrapped along their lengths. Several lines of evidence indicate that neuronal activity can influence myelination, however, the cellular and molecular mechanisms that mediate communication between axons and oligodendrocytes remain poorly understood. Prior research showed that the neurotrophic growth factor Bdnf and its receptor Ntrk2 promote myelination in rodents, raising the possibility that Bdnf and Ntrk2 convey myelin-promoting signals from neurons to oligodendrocytes. We explored this possibility using a combination of gene expression analyses, gene function tests, and myelin sheath formation assays in zebrafish larvae. Altogether, our data indicate that, although not essential for myelination, Bdnf-Ntrk2 signaling contributes to the timely formation of myelin in the developing zebrafish spinal cord.

## INTRODUCTION

Myelin internodes that form on central nervous system (CNS) axons can vary in thickness and length and in the total amount of axon coverage (Almeida et al., 2011a; Chong et al., 2012; Murtie et al., 2007; Tomassy et al., 2014). Myelin characteristics on individual axons can differ, at least in part, according to neuronal subtype (Nelson et al., 2020; Tomassy et al., 2014; Zonouzi et al., 2019), suggesting that axon-specific cues differentially regulate myelination. Additionally, myelin characteristics can change in response to neuronal activity (Bacmeister et al., 2022; Chorghay et al., 2022; Hines et al., 2015; Mensch et al., 2015; Mitew et al., 2018), consistent with the possibility that axons provide cues to oligodendrocytes (OLs) that regulate myelin sheath formation. Axons secrete vesicle contents at points of contact with nascent myelin sheaths (Almeida et al., 2021; Hughes and Appel, 2019) and vesicle secretion promotes myelin sheath growth and stability (Almeida et al., 2021; Hines et al., 2015). Thus, activity-evoked axon vesicle secretion could serve as one mechanism contributing to myelin formation by presenting axon subtype-specific pro-myelinating factors to OLs.

One potential axon secreted pro-myelinating molecule is the growth factor Bdnf. *Bdnf* homozygous mutant mice had a deficit of myelinated optic nerve axons and expressed abnormally low levels of *Mbp* and *Plp* mRNA in the cortex and hippocampus (Cellerino et al., 1997), and low levels of Mbp detected by immunohistochemistry and Western blot (Djalali et al., 2005). Both homozygous and heterozygous mutant mice had fewer NG2^+^ oligodendrocyte precursors cells (OPCs) in the basal forebrain than wild type, and although *Bdnf*^+/–^ mice had normal numbers of CC1^+^ oligodendrocytes, the intensity of Mbp immunostaining was reduced (Vondran et al., 2010). *Bdnf*^+/–^ mice were hypomyelinated throughout the brain and spinal cord, and application of Bdnf to neuron and OL co-cultures promoted myelination (Xiao et al., 2010). These observations indicate that Bdnf positively regulates oligodendrocyte lineage cell (OLC) number and myelination. By contrast, in cell culture proBdnf inhibited neural stem cell proliferation and reduced formation of neurons and glia, including OLs (Li et al., 2017). The receptor for Bdnf, Neurotrophic tyrosine kinase receptor 2 (Ntrk2, also known as NtrkB and TrkB) is also implicated in myelination, as OL expression of the receptor mediates remyelination following injury (Fletcher et al., 2018). Whether Bdnf acts as a direct axon to OLC signal to regulate OL differentiation and myelin sheath characteristics is not known.

We designed this study to investigate the possibility that Bdnf functions as an axon to OLC promyelinating factor. As a first step we investigated *bdnf* and *ntrk2* expression and assessed oligodendrocyte development and myelination in zebrafish larvae lacking *Bdnf* and *Ntrk2* functions to determine the effect of Bdnf-Ntrk2 signaling on developmental myelination. These experiments revealed subtle, transient alterations of OL development and myelin sheath formation resulting from loss of Bdnf-Ntrk2 signaling, supporting prior data from rodent systems indicating that Bdnf and Ntrk2 contribute to developmental myelination.

## RESULTS

### Oligodendrocyte lineage cells express *ntrk2a* but little *ntrk2b* or *bdnf* during development

As a first step toward testing whether Bdnf-Ntrk2 signaling regulates developmental myelination in zebrafish, we examined transcript levels of *bdnf* and two *ntrk2* paralogs, *ntrk2a* and *ntrk2b*, in RNA-seq data obtained from transgenically marked OPCs and OLs that were sorted from 7 days postfertilization (dpf) zebrafish larvae (Ravanelli et al., 2018). These data revealed that OPCs expressed substantial levels of *ntrk2a* whereas OLs expressed *ntrk2a* at a lower level (Figure 1A). By contrast, neither OPCs nor OLs expressed appreciable levels of *ntrk2b* or *bdnf* (Figure 1A). Because OPCs initially arise at about 40 hours postfertilization (hpf) and myelination begins at about 3 dpf in zebrafish, we next used a hybridization chain reaction (HCR) method to detect transcripts in situ across earlier developmental stages. At 2 dpf numerous spinal cord cells expressed *ntrk2a*, but little expression was evident in cells that expressed *sox10*, which marks oligodendrocyte lineage cells (OLCs) consisting of both OPCs and OLs (Figure 1B). At 3 dpf many spinal cord cells, including *sox10*^+^ OLCs, appeared to uniformly express *ntrk2a* (Figure 1C). At 5 dpf, a period of active myelination, *sox10*^+^ OLCs continued to express *ntrk2a* (Figure 1D). By contrast, we detected little spinal cord expression of *ntrk2b* transcripts across these developmental stages (Figure 1E and data not shown). These data indicate that OLCs express *ntrk2a* but little detectable *ntrk2b*.

**Figure 1.**
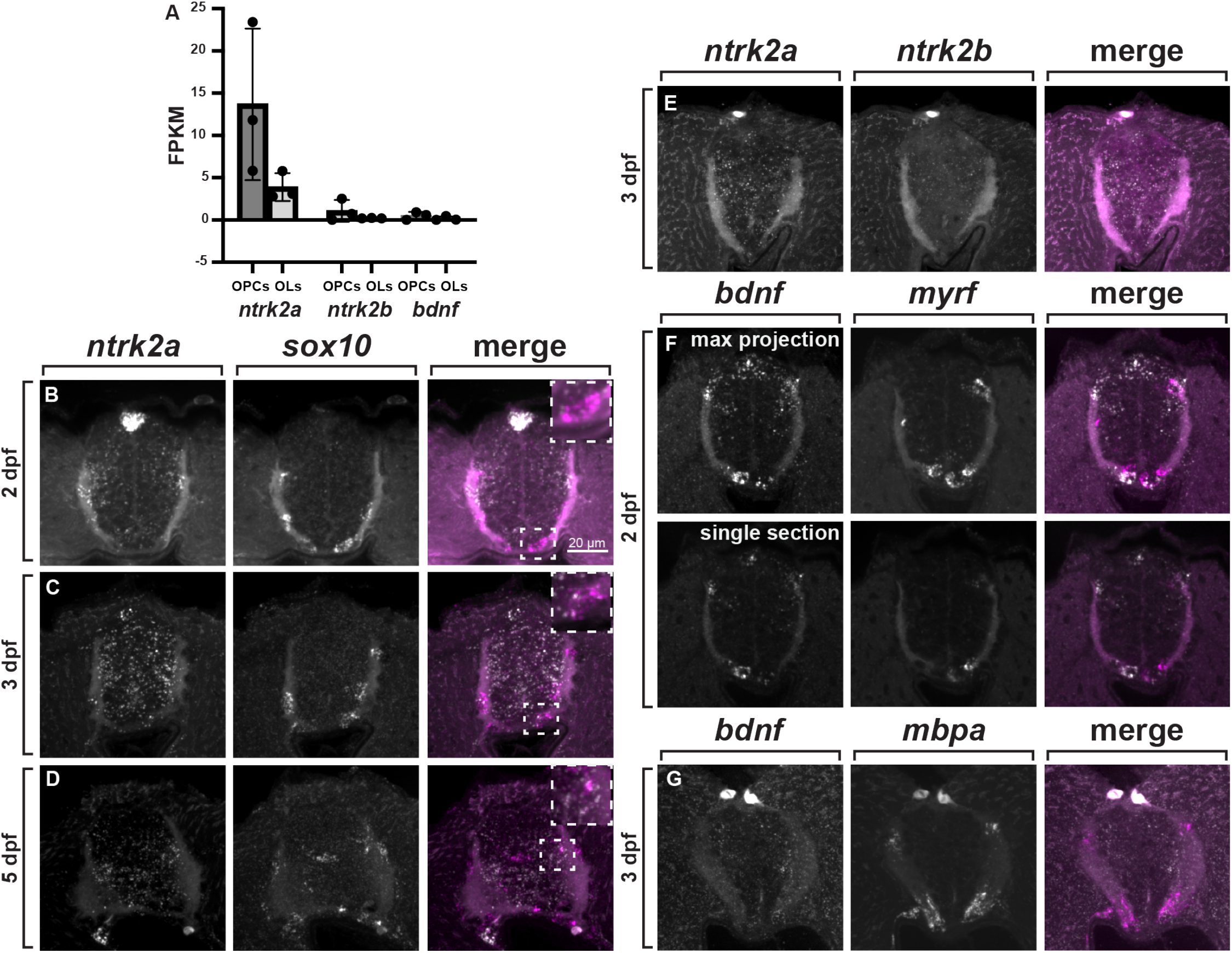
OLCs express *ntrk2a* but not *ntk2b* or *bdnf*. (A) Expression (FPKM) of *ntrk2a, ntrk2b*, and *bdnf* in OPCs and OLs isolated from 7 dpf larvae. (B-G) Representative transverse sections, dorsal up, of trunk spinal cords processed for in situ RNA hybridization. *sox10*^*+*^ OPCs express little detectable *ntrk2a* RNA at 2 dpf (B) but do express *ntrk2a* at 3 and 5 dpf (C, D). Insets show enlargements of outlined regions. At 3 dpf, spinal cord cells express little *ntrk2b* (E). At 2 dpf, cells close to *myrf*^+^ pre-myelinating OLs express *bdnf* (F) but little *bdnf* expression is evident near *mbpa*^*+*^ OLs at 3 dpf (G).

At 2 dpf, *bdnf* transcripts appeared to be concentrated in dorsal and ventral spinal cord, close to but not including cells that expressed *myrf*, which marks OPCs and pre-myelinating and myelinating OLs (Figure 1F). At 3 and 5 dpf *bdnf* RNA appeared to be less abundant and more uniformly distributed throughout the spinal cord (Figure 1G and data not shown). These observations raise the possibility that cells located close to OLCs are a source of Bdnf during a period of OPC to OL differentiation. Altogether, these data are consistent with a possibility that Bdnf-Ntrk2 signaling promotes developmental myelination.

### *bdnf* and *ntrk2a* regulate OLC number and differentiation during development

To test the possibility that Bdnf–Ntrk2 signaling promotes developmental myelination in zebrafish, we used CRISPR/Cas9 gene targeting to create loss-of-function mutations of *bdnf, ntrk2a*, and *ntrk2b* (Figure 2A). Each of the mutations that we selected are nonsense alleles, therefore, the mutant transcripts are potentially subject to nonsense mediated decay (NMD). Because NMD can cause transcriptional adaptation of related genes (El-Brolosy et al., 2019), *ntk2a* and *ntrk2b* encode similar proteins, and spinal cords of wild-type embryos express *ntrk2a* but little *ntrk2b*, we used HCR to detect *ntk2a* and *ntrk2b* transcripts expressed by *ntk2a* mutant larvae. At 3 dpf, *ntk2a*^*–/–*^ larvae had little detectable *ntrk2a* transcripts (Figure 2B,C), suggesting that RNA encoded by the *co7005* allele is subject to NMD. At 5 dpf, we could detect *ntrk2a* transcripts in mutant larvae at levels comparable to or greater than in wild-type larvae (Figure 2D,E). At both 3 and 5 dpf, *ntk2a*^*–/–*^ larvae expressed considerably more *ntrk2b* RNA than comparable staged wild-type larvae (Figure 2F-I). Thus, NMD of *ntrk2a* transcripts might trigger a transcriptional adaptation response that elevates *ntrk2b* expression.

**Figure 2.**
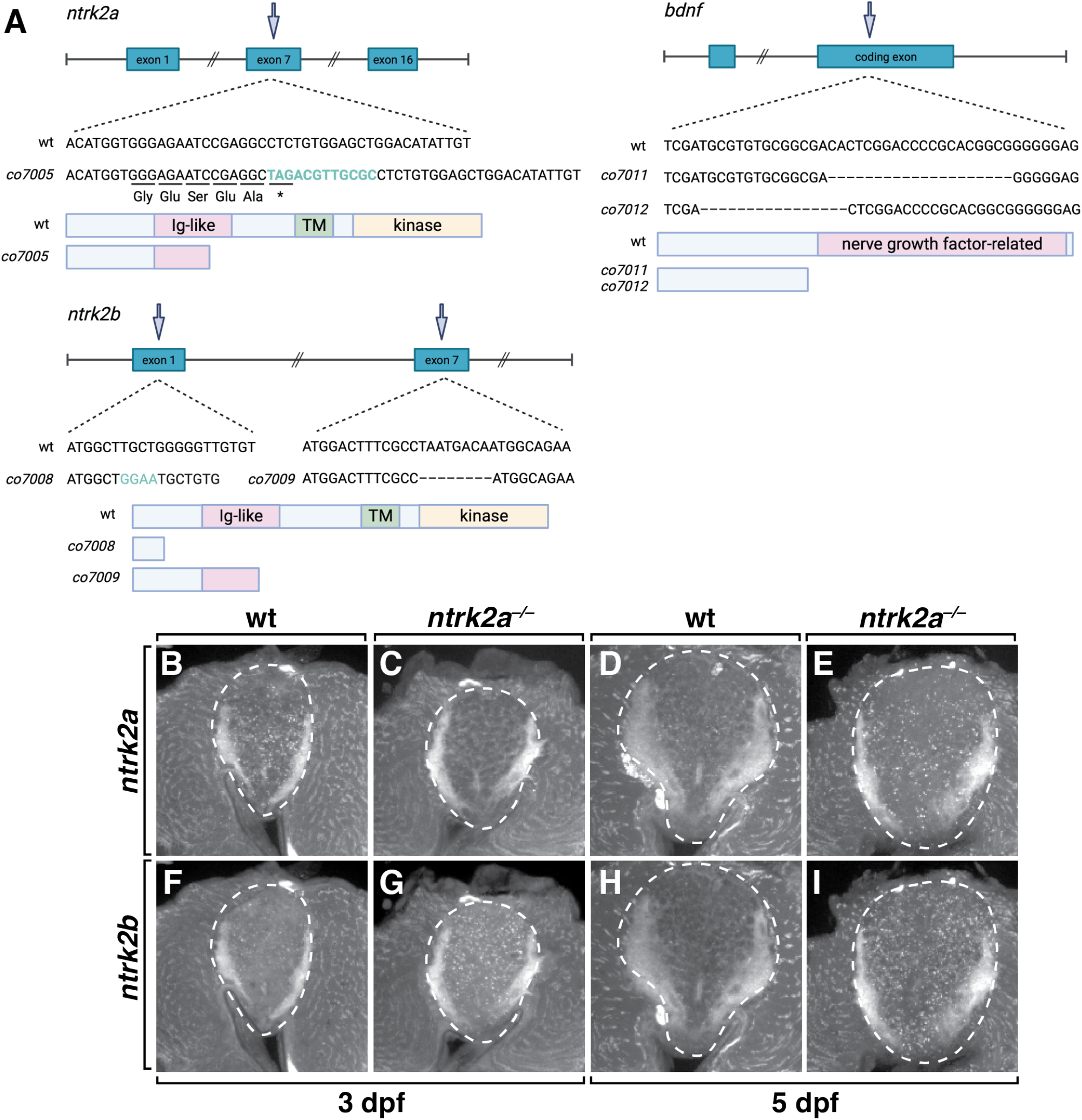
*ntrk2* and *bdnf* alleles and potential *ntrk2* transcriptional adaptation. (A). Schematic representations of *ntrk2a, ntrk2b*, and *bdnf* mutant alleles and predicted truncated proteins. Arrows mark positions of mutant alleles. Wild-type and mutant DNA sequences are shown with corresponding wild-type and mutant protein structures. (B-I) Representative transverse sections of trunk spinal cord, dorsal up, processed for in situ RNA hybridization. At 3 dpf, *ntrk2a* transcripts are absent from *ntrk2a*^*co7005*^ homozygous mutant larva but present in wildtype, suggesting that *ntrk2a* mRNA is cleared by nonsense mediated decay (B, C). At 5 dpf, *ntrk2a* homozgous mutant spinal cord expresses more *ntrk2a* transcripts than wildtype (D, E). *ntrk2a* homozgous mutant larvae express higher levels of *ntrk2b* transcripts than wild-type larvae at 3 and 5 dpf, suggestive of transcriptional adaptation (F-I). Dashed lines outline the spinal cord.

Having established mutant lines we analyzed the number and distribution of OLCs in homozygous mutant larvae compared to wild-type siblings. Because OLCs do not express *ntrk2b* at levels detected by either RNA-seq or in situ RNA hybridization we initially limited our analyses to *ntrk2a* and *bdnf* mutant larvae. For all the following data we used the *ntrk2a*^*co7005*^ allele and the *bdnf*^*co7011*^ allele. At 3 dpf, OLCs express *olig2* and *sox10*. Therefore, we labeled tissue sections obtained from wild-type and mutant larvae carrying a *Tg(olig2:EGFP)* transgene (Shin et al., 2003) with an anti-Sox10 antibody (Park et al., 2005). In wild-type larvae, OPCs and myelinating OLs occupy ventral and dorsal spinal cord, corresponding to longitudinal myelinated axons tracts, and a few OPCs occupy the neuron-rich region of the medial spinal cord (Fig. 3A). The distribution of OLCs in *bdnf*^*–/–*^ and *ntrk2a*^*–/–*^ larvae appeared to be similar to wild-type larvae (Fig. 3A-C), indicating that loss of these functions did not appreciably affect OPC migration. Notably, *bdnf* mutant larvae had a small but statistically significant excess of OLCs compared to their wild-type siblings (Fig. 3B,E). However, OLC number was the same in *ntrk2a* mutant larvae and their wild-type siblings (Fig. 3C,F). Because these data indicate that *bdnf*^*–/–*^ larvae but not *ntrk2a*^*–/–*^ larvae have a small excess of OLCs, we reasoned that the elevated expression of *ntrk2b* that we detected in *ntrk2a*^*–/–*^ larvae could suppress a phenotype resulting from loss of Ntrk2a function. We therefore intercrossed *ntrk2a*^*+/–*^;*ntrk2b*^*+/–*^ adults to produce double homozygous mutant larvae. Labeling tissue sections with anti-Sox10 antibody revealed no difference in OLC number between wild-type and *ntrk2a*^*–/–*^;*ntrk2b*^*+/–*^ larvae (Fig. 3D,G). These data suggest that Bdnf appears to limit the number of OLCs formed in the zebrafish spinal cord, but not their specification or distribution, and that the effect of Bdnf on OLC number is not mediated by Ntrk2 receptors.

**Figure 3.**
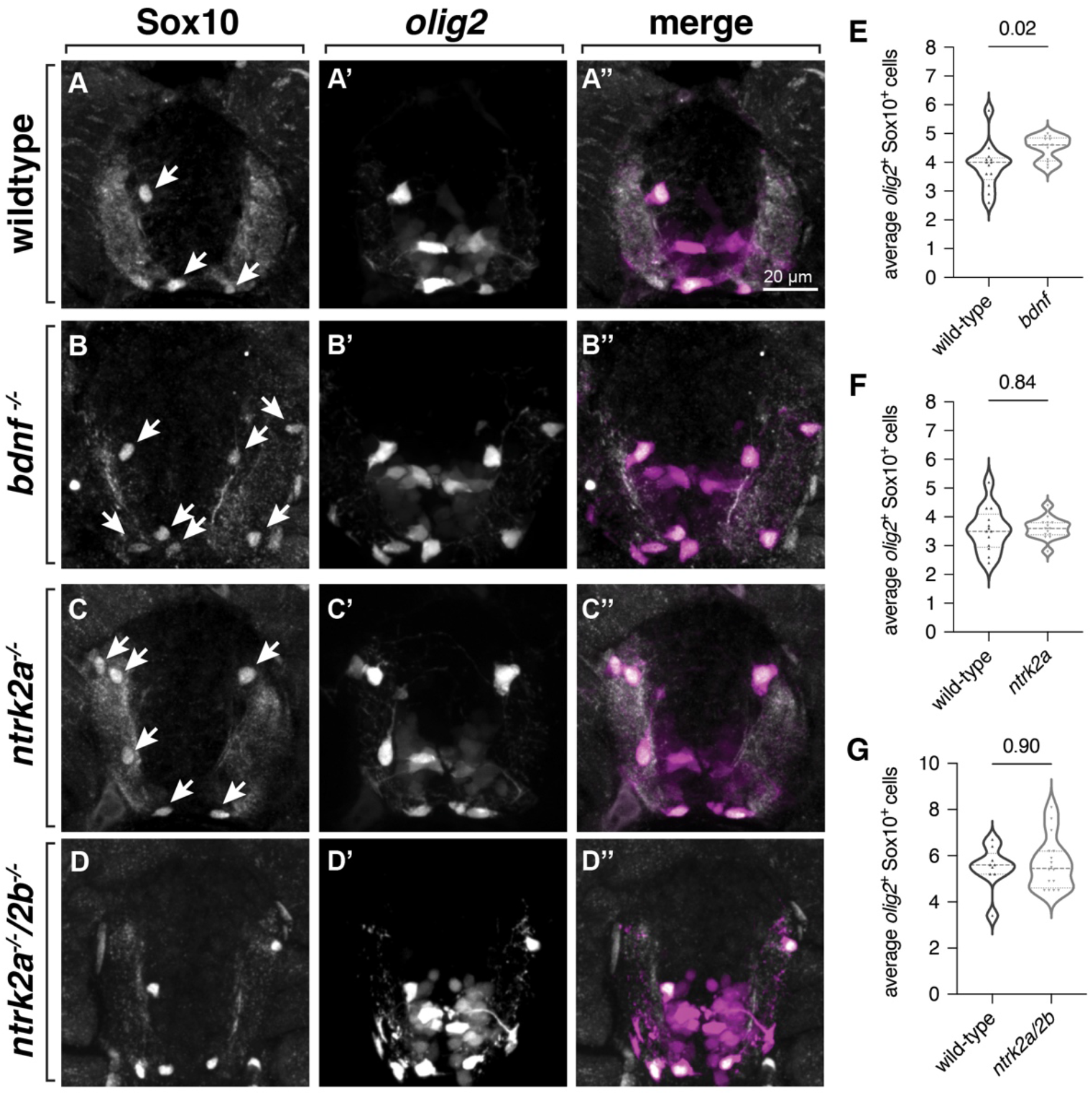
Loss of *bdnf* but not *ntrk2a* alters OLC number. (A-D) Representative images of transverse sections at the level of trunk spinal cord with dorsal up, processed to detect Sox10 protein. All larvae are 3 dpf and have the *Tg(olig2:EGFP)* transgene. Arrows mark Sox10^+^ *olig2:*EGFP^+^ OLCs. *Bdnf* homozygous mutant larve have a small but significant excess of OLCs compared to wildtype (A, B, E) whereas the number of OLCs in *ntrk2a* and *ntrk2a;ntrk2b* homozgous mutant larvae is similar to wildtype (C, D, F, G). Sample sizes: n^WT^=13 larvae, n^*bdnf*^=13; n^WT^=13, n^*ntrk2a*^=14; n^WT^=9, n^*ntrk2a/2b*^=16. Significance determined by unpaired t-tests (*bdnf, ntrk2a*) and a Mann-Whitney test (*ntrk2a/ntrk2b*).

To investigate the possibility that Bdnf-Ntrk2 signaling regulates OLC differentiation, we next performed in situ hybridization to detect expression of *mbpa* RNA, which encodes Myelin basic protein (Mbp), a marker of mature OLs. Despite having no difference in spinal cord OLC number compared to wild type, *ntrk2a*^*–/–*^ larvae had a small excess of *mbpa*^+^ cells (Fig. 4A,B,D). Additionally, *mbpa*^+^ cells occasionally occupied the neuron-dense portion of the spinal cord (Fig. 4B), which, in wild-type larvae, is typically occupied by OPCs but not OLs. By contrast, *bdnf*^*–/–*^ larvae had the same number of *mbpa*^+^ OLs as their wild-type siblings (Fig. 4A,B,D), even though they had an excess of spinal cord OLCs. As an additional measure of differentiation we also quantified the number of *mbpa*^+^ puncta within each cell. This revealed that whereas the amount of *mbpa* transcripts expressed by *ntrk2a*^*–/–*^ larvae was less than their wild-type siblings, the level of *mbpa* expression by *bdnf*^*–/–*^ larvae was unchanged (Fig. 4F,G). These observations raise the possibility that Ntrk2 has at least two functions in OL differentiation. First, the excess number and ectopic position of OLs in *ntrk2a*^*–/–*^ larvae is consistent with a possibility that Ntrk2 function prevents premature OL differentiation. Second, the reduced level of *mbpa* transcripts in mutant larvae suggest that Ntrk2 also promotes OL differentiation. Because we found no differences in these measurements in *bdnf*^*–/–*^ larvae, our results also raise the possibility that Ntrk2 contributes to OL differentiation independently of Bdnf function.

**Figure 4.**
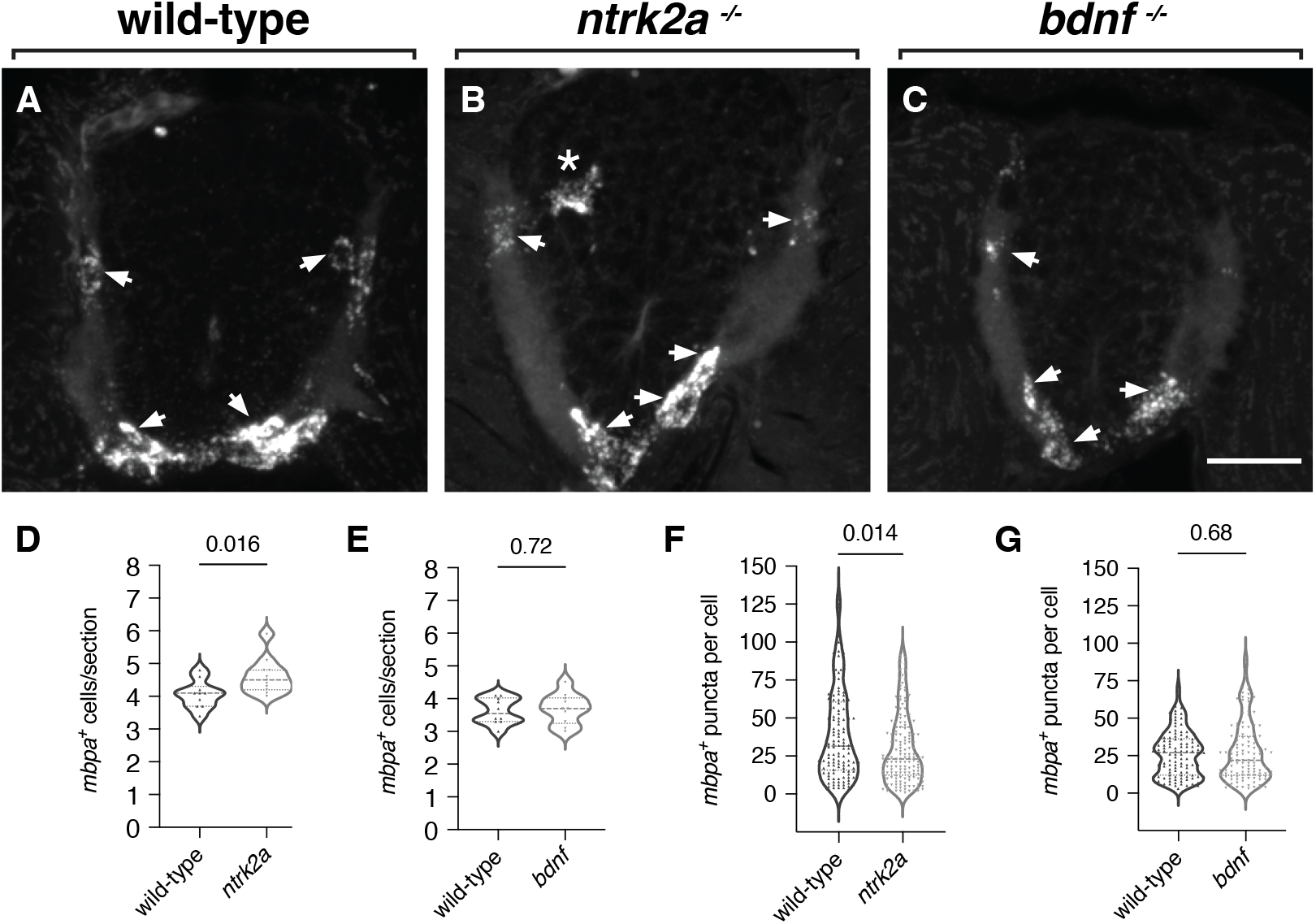
Loss of *ntrk2a* but not *bdnf* function alters the number of *mbpa*^+^ OLs and the level of *mbpa* expression. (A-C) Representative transverse sections of 3 dpf larvae at the level of the trunk spinal cord processed for in situ RNA hybridization to detect *mbpa* transcripts (*mbpa*^*+*^ OLs denoted with arrowheads). *ntrk2a* homozygous mutant larvae occasionally have ectopically-positioned *mbpa*^*+*^ OLs (B, asterisk). *ntrk2a* mutant larvae have more *mbpa*^*+*^ OLs than wildtype (D) whereas the number of *mbpa*^*+*^ OLs is not different from wildtype in *bdnf* mutant larvae (E). *ntrk2a* mutant larvae have fewer *mbpa*^*+*^ puncta per cell than wildtype (F) whereas wild-type and *bdnf* mutant larvae have similar numbers of puncta (G). Scale bar = 20 um. n^WT^=10 larvae, n^*ntrk2a*^=11; n^WT^=10, n^*bdnf*^=10. Statistical significance determined by an unpaired t-test (*bdnf* cell counts) and Mann-Whitney tests (*ntrk2a* cell counts, puncta counts).

### Ntrk2 regulates initial myelination, but Bdnf is dispensable

Changes in OL production and differentiation in *ntrk2a* mutants could in turn influence myelin sheath formation by OLs. Myelin sheaths differ in number and length and both attributes are sensitive to perturbations that influence myelination. To test this possibility, we measured the number and length of myelin sheaths formed by individual OLs in both *ntrk2a* and *bdnf* mutant larvae. To perform this experiment we used the Tol2 transgenesis system and microinjection to express membrane-tethered EGFP under control of *mbpa* regulatory DNA in larvae. This approach labels a subset of OLs allowing us to accurately measure sheath length and number. We quantified sheath characteristics in *nrtk2a* and *bdnf* mutants, and wild-type controls at 3 dpf, when myelination initiates, and 5 dpf, when sheaths are more stable. At 3 dpf, we found that myelin sheaths were dramatically shorter in *ntrk2a* mutants compared to wild-type controls (a ~34% reduction), with no effect on the average number of sheaths generated (Figure 5A,C,E,G). In contrast, we found no change in sheath characteristics in *bdnf* mutants compared to controls (Figure 5B,D,F). At 5 dpf, neither sheath length or number differed in *bdnf* and *ntrk2a* mutant larvae compared to wild-type siblings (Figure 5H-N). We conclude from these data that Ntrk2a and not Bdnf regulates early myelination in the spinal cord, but hypomyelination in *ntrk2a* mutants does not persist.

**Figure 5.**
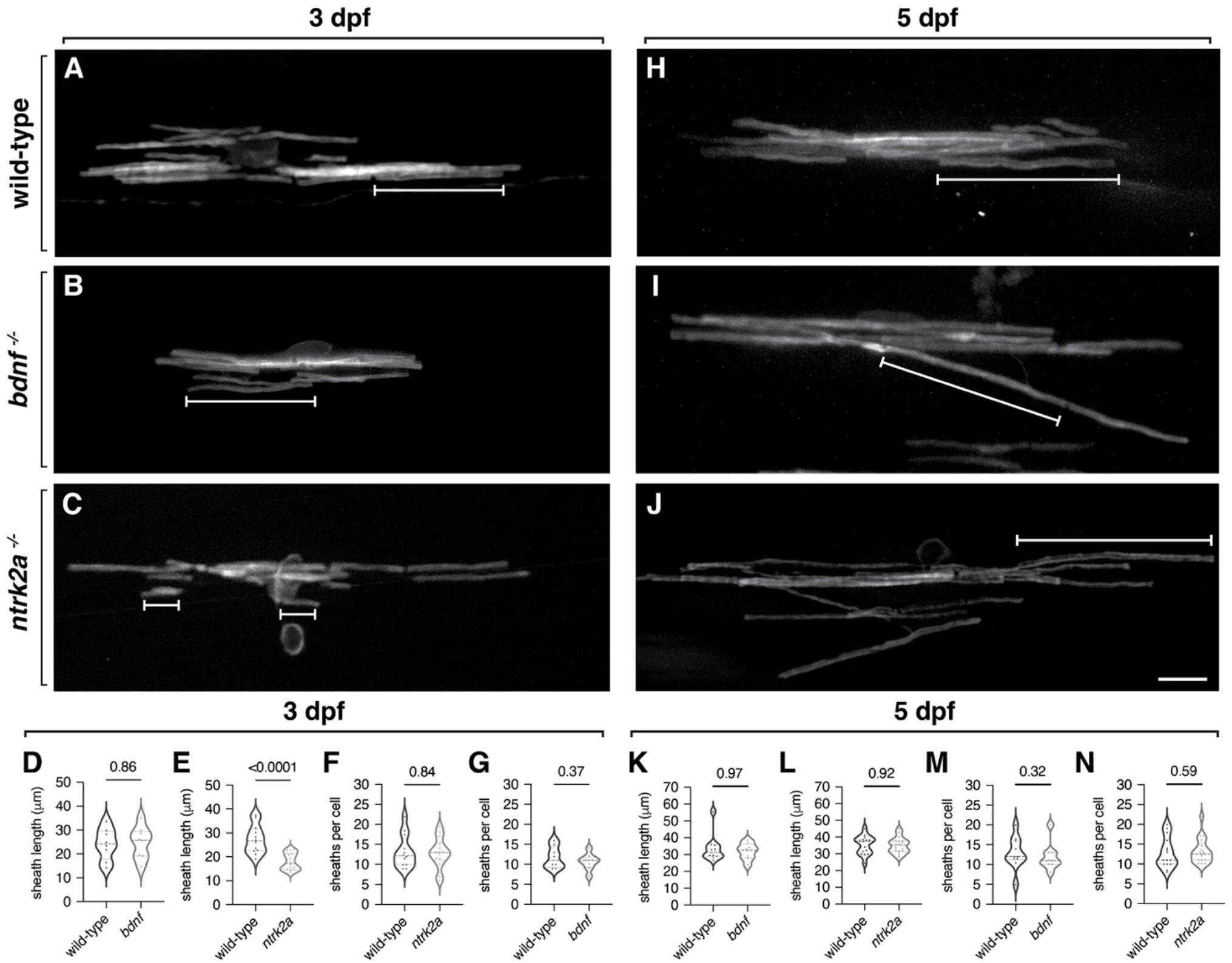
Loss of *ntrk2a* function leads to a stage-specific reduction in myelination. Representative images of individual oligodendrocytes in living 3 dpf (A-C) and 5 dpf (H-J) larvae expressing *mbpa:*EGFP-CAAX. Brackets indicates myelin sheaths. At 3 dpf, the average and cumulative length of myelin sheaths were reduced in *ntrk2a* mutants compared to wild-type, while myelin characteristics in *bdnf* mutants were not significantly different than wildtype (E-G). At 5 dpf, the number and average length of myelin sheaths formed by oligodendrocytes is similar in wild-type and mutant larvae (K-N). 3 dpf: n^WT^= 12 larvae, n^*bdnf*^ = 8; n^WT^= 8, n^*ntrk2a*^ = 14. 5 dpf: n^WT^= 10, n^*bdnf*^ = 8; n^WT^ = 15, n^*ntrk2a*^ = 17. Statistical analysis performed using unpaired t-tests (3 dpf datasets) and Mann-Whitney tests (5 dpf datasets). Scale bar = 10 um.

## DISCUSSION

Conceptually, growth factor-mediated intercellular signaling seems well suited to convey promyelinating signals from axons to nascent myelin sheaths. Consistent with this view, prior research has provided evidence that some growth factor signaling pathways are important drivers of myelination. For example, elimination of FGF receptor 2 (FGFR2) from oligodendrocytes reduced myelin sheath thickness (Furusho et al., 2017, 2012). FGFR2 localizes to myelin sheath paranodes, positioning it to possibly receive FGF secreted by axons (Furusho et al., 2017). Additionally, ciliary neurotrophic factor (CNTF) promoted myelination of oligodendrocytes in cell culture (Stankoff et al., 2002). Dense core vesicles at excitatory presynaptic terminals contain Bdnf (Dieni et al., 2012), potentially representing a source of neuronal, promyelinating signal. However, astrocytes and microglia also express growth factors and other molecules that have promyelinating functions (Traiffort et al., 2020). The exact sources of promyelinating factors and how they are received by oligodendrocytes to regulate myelination remain poorly understood.

Based on evidence from rodents that Bdnf-Ntrk2 signaling promotes myelination (Cellerino et al., 1997; Djalali et al., 2005; Fletcher et al., 2018; Vondran et al., 2010; Xiao, 2023; Xiao et al., 2010), we examined OLC populations and myelin sheaths in *bdnf* and *nrtk2a* mutant zebrafish larvae. Regarding *bdnf* mutants, we found an increase in total OLCs but no changes in mature OLs or myelin sheaths. In contrast, we found an increased number of OLs in *ntrk2a* mutants, but these OLs contained fewer *mbpa* transcripts than OLs in control larvae. This may suggest Ntrk2a regulates OL differentiation or maturation. In line with this, we found a dramatic reduction in myelin sheath length in *ntrk2a* mutants at the initial stage of myelination. Although this phenotype did not persist into later stages, these early myelin deficits potentially influence circuit formation and function.

In both mouse and zebrafish, the OLC and myelin phenotypes resulting from loss of Bdnf-Ntrk2 signaling are subtle and transient. Thus, Bdnf and Ntrk2 functions are not essential for myelination, but yet they contribute to the timely production of myelin. Notably, OLs do not require biological signals for myelination because they can myelinate fixed axons and synthetic fibers (Bechler et al., 2015; Lee et al., 2012; Rosenberg et al., 2008). Furthermore, myelin sheath length is an inherent property of OLs, reflecting distinct CNS origins (Bechler et al., 2015). However, the number of OLCs and myelin sheath properties such as number, length, and thickness, can be modified, particularly by neuronal activity (Hines et al., 2015; Mensch et al., 2015; Mitew et al., 2018). One interpretation of these observations is that there are two features of developmental myelination consisting of an intrinsic capacity to myelinate that can be modulated by extrinsic factors as a mechanism of adaptation (Bechler et al., 2018). Potentially, Bdnf is one of numerous extrinsic factors that act on OLCs to modulate myelin production.

Bdnf is a particularly intriguing candidate for regulating myelination because of its well-established role in synaptic plasticity. The *Bdnf* locus has multiple alternative transcription start sites, each of which produces a distinct mRNA isoform containing a common exon that encodes BDNF (Aid et al., 2007; Timmusk et al., 1993). A subset of mRNA isoforms, which were differentially responsive to kainic acid-induced seizures, was detected in rodent brains (Aid et al., 2007; Metsis et al., 1993; Timmusk et al., 1993) suggesting that they are transcriptionally regulated by neuronal activity. Consistent with this, expression of exon III-containing mRNA was induced by membrane depolarization via calcium signaling (Shieh et al., 1998). Results from co-culture experiments indicated that Bdnf can promote activity-regulated myelination (Lundgaard et al., 2013). More recently, activity-regulated expression of Bdnf was shown to promote OPC proliferation and myelination of cortical projection neurons in adult mice (Geraghty et al., 2019). These observations raise the possibility that, during development, Bdnf, signaling through Ntrk2 receptors, influences the amount of myelin on specific axons in response to neural circuit activity.

There are several important limitations to our investigation. First, the mutations we used eliminate Bdnf and Ntrk2 functions in all cells of developing larvae. Consequently, we currently are not able to discriminate between functions in different neural cell types that might be important for myelination. Second, we do not have a way of visualizing Bdnf in vivo, therefore we are unable to learn if Bdnf might be delivered to OLCs from axons or astrocytes or some other source. Third, we have not yet addressed the possibility that Bdnf mediates promyelinating effects of of neuronal activity on myelination. Doing this will require combining methods for Bdnf visualization, cell type-specific gene inactivation, and chemogenetic or genetic manipulation of neuronal activity.

In summary, our data support conclusions drawn from studies using rodents that Bdnf-Ntrk2 signaling contributes to developmental myelination, but that this signaling pathway is likely only one of many that determine the myelin landscape.

## MATERIALS AND METHODS

### Zebrafish lines and husbandry

The Institutional Animal Care and Use Committee at the University of Colorado School of Medicine approved all animal work, which follows the US National Research Council’s Guide for the Care and Use of Laboratory Animals, the US Public Health Service’s Policy on Humane Care and Use of Laboratory Animals, and Guide for the Care and Use of Laboratory Animals. Larvae were raised at 28.5°C in embryo medium and staged as hours (hpf) according to morphological criteria (Kimmel et al., 1995). Zebrafish lines used in this study included AB wildtype and *Tg(olig2:EGFP)* (Shin et al., 2003). Because zebrafish sex determination does not occur until juvenile stages, we were unable to determine the sex of the embryos and larvae in our experiments.

### CRISPR/Cas9 mutagenesis and genotyping

For generation of *bdnf, ntrk2a*, and *ntrk2b* loss-of-function mutants, a CRISPR-Cas9 mutagenesis strategy involving injection of single guide RNAs (sgRNAs) and Cas9 mRNA was employed. Genomic targets for Cas9 nuclease were identified using ZiFIT Targeter Software (Sander et al., 2010) and searched for potential ‘off-target’ effects using BLASTN (National Center for Biotechnology Information). Single stranded DNA oligonucleotides targeting the genomic targets were synthesized (Table 1), annealed and inserted into the pDR274 vector to generate a plasmid for synthesis of sgRNAs as described previously (Jewett et al., 2020). Cas9 mRNA was transcribed using the pT3Ts vector (Addgene). The sgRNA (125 ng/nl) and Cas9 mRNA (100 ng/nl) were coinjected into 1-cell stage embryos in a solution containing 100 mM KCl and 0.2% phenol red. Adults were genotyped by sequencing the targeted genomic regions via polymerase chain reaction using primers that flanked the targeted site. Genotyping was performed using the PCR primers listed in Table 1.

**Table 1.**
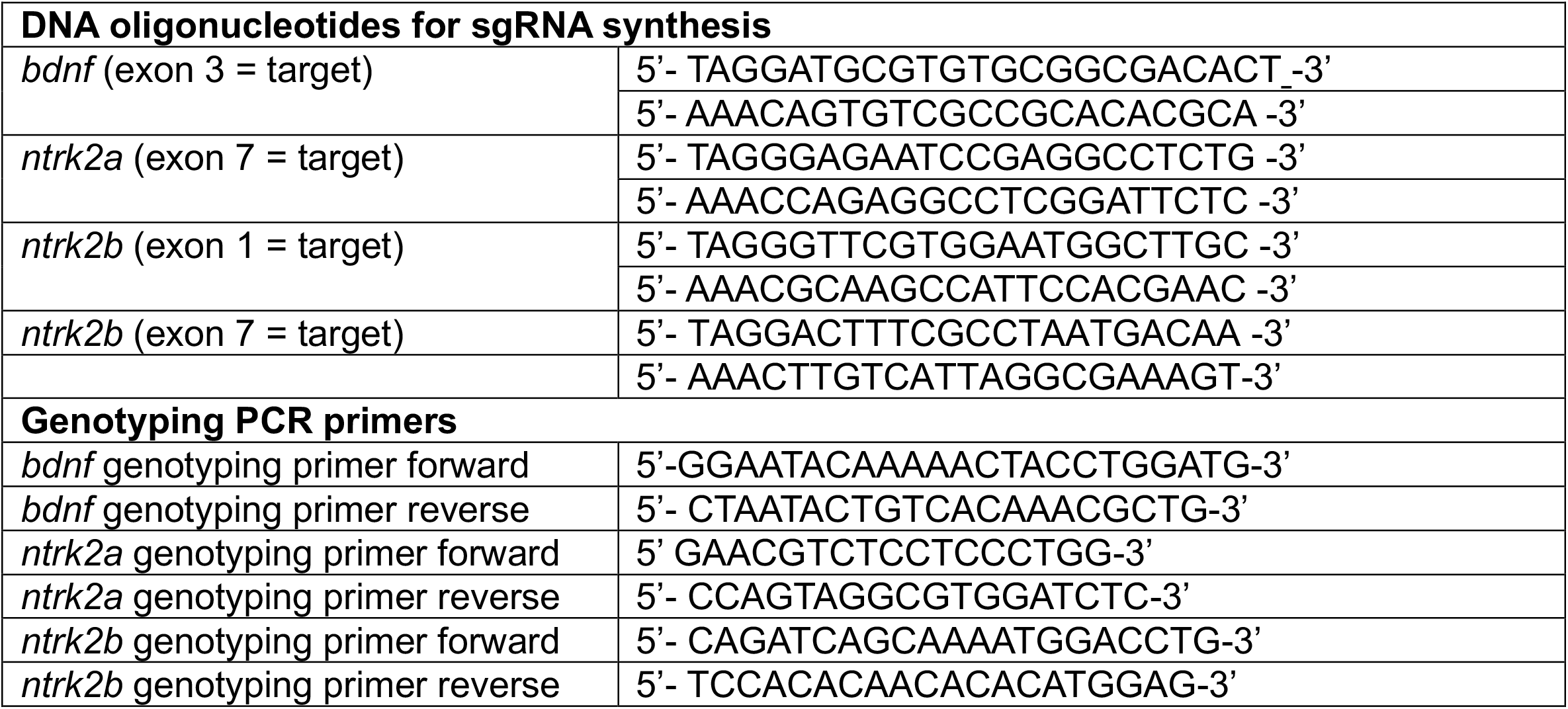

### Immunohistochemistry

Larvae were anesthetized using Tricaine and fixed using 4% paraformaldehyde/1X PBS overnight at 4°C. Embryos were washed with 1X PBS, rocking at room temperature, and embedded in 1.5% agar/5% sucrose, frozen over dry ice and sectioned in 15 μm transverse increments using a cryostat microtome. Slides were placed in Sequenza racks (Thermo Scientific), washed 3×5 min in 0.1%Triton-X 100/1X PBS (PBSTx), blocked 1 hour in 2% goat serum/2% bovine serum albumin/PBSTx and then placed in primary antibody (in block) overnight at 4°C. The primary antibodies used was rabbit anti-Sox10 (1:500) (Park et al., 2005). Sections were washed for 1 hr at room temperature with PBSTx and then incubated for 1 hr at room temperature with secondary antibodies at a 1:200 dilution in block. The secondary antibody used was AlexaFluor 546 anti-rabbit (Invitrogen A-11010). Sections were washed for 1 hr with PBSTx and mounted in VectaShield (Vector Laboratories). Sox10^+^*olig2*:EGFP^+^ cells were counted and averaged for 10 sequential sections for each embryo genotype and statistical analysis obtained using Prism software, version 10.4.0

### In situ RNA hybridization

The following probes were synthesized by Molecular Instruments: *ntrk2a* (B1), *ntrk2b* (B2), *bdnf* (B3), and *mbpa* (B3) for use with B1-B3 488, 546, and 647 fluorophores. Fluorescent in situ hybridization (FISH) procedure was guided by the in situ hybridization chain reaction protocol for whole-mount zebrafish (Molecular Instruments v3.0; Choi et al., 2018). Larvae were euthanized using Tricaine at given timepoints and fixed using 1 mL 4% paraformaldehyde/1xPBS overnight at 4°C. The following day, larvae were rinsed with 1 mL 1x PBS and transferred to 1 mL 100% MeOH and stored at −20°C overnight prior to use. Larvae were rehydrated with a series of graded 1 mL MeOH/PSBTw washes (1x 75% MeOH, 1 × 50% MeOH, 1x 25% MeOH, and 5 × 100% PBST) for 5 min each at room temperature. Tissue was permeabilized with proteinase K (2 dpf at 1:200 for 8 min, 3 dpf at 1:200 for 12 min, and 5 dpf at 1:1000 for 40 min) and immediately washed two times with 1 mL PBSTw. Samples were post-fixed with 1 mL 4% paraformaldehyde/1x PBS for 20 min at room temperature on a rocker, followed by 5 × 5-minute washes with 1mL PBST. For the detection stage, 500 uL pre-warmed probe hybridization buffer was added and incubated for 30 min at 37°C. Then, the pre-hybridization solution was removed and the prewarmed probe solution was added (2 pmol total: 2 uL of probe/500 uL probe hybridization buffer) and samples were incubated overnight (12-16 hours) at 37°C. The next day, probe wash buffer was prewarmed to 37°C before starting the wash step.

Samples were washed 4 × 1 5 min with 500 uL probe wash buffer at 37°C, followed by two 5 min washes with 1 mL 1 X saline sodium citrate (SSC), with 0.1% Tween (SSCT) at room temperature. Before the last probe wash buffer wash, the amplification buffer was equilibrated to room temperature. For the amplification stage, samples were pre-amplified with 500 uL room temperature amplification buffer for 30 min at room temperature. During the pre-amplification step, 30 pmol of hairpin h1 and 30 pmol of hairpin h2 were separately prepared by snap cooling 10 uL of 3 uM hairpin stock at 95°C for 90 seconds and were allowed to cool to room temperature in a dark drawer for 30 min prior to use. The hairpin solution was then prepared by adding the snap-cooled hairpins to 500 uL amplification buffer at room temperature. After completing the pre-amplification step, the pre-amplification solution was removed, 500 uL hairpin solution was added to each tube, and samples were incubated in a dark drawer at room temperature overnight (12-16 hours). The following day, the excess hairpins were removed by a series of washes of 500 uL 5 x SSCT for 2 × 5 min, 2 × 30 min, and 1 × 5 min. Samples were post-fixed with 1 mL of 4% paraformaldehyde/1xPBS for 20 min at room temperature on a rocker. The samples were washed 2 x with 1 mL of 1 x PBS diethyl pyrocarbonate (DEPC) with no incubation time. Larvae were embedded in 1.5% agar/5% sucrose, frozen over dry ice and sectioned in 15 μm transverse increments using a cryostat microtome. Slides were mounted with vectashield. *mbpa*^+^ cells were counted and averaged for 10 sequential sections for each embryo genotype and statistical analysis obtained using Prism software, version 10.4.0.

Using Imaris software, version 10.2.0, images were deconvoluted and processed in Surfaces. *Mbpa* 546 hairpin fluorophore ROIs were selected, thresholding set to 0.8 um and filter seed added to remove background. Puncta within 3 sequential sections for each of 10 *bdnf* mutant and wild-type siblings and 10 *ntrk2a* mutant and wild-type were collected for non-overlapping cells, analyzed and statistical analysis obtained using Prism Software, version 10.4.0.

### Myelin sheath analysis

To analyze myelin sheath length and number of we injected approximately 3 nl of the plasmid *pEXPRS*.*mbpa:EGFP-CAAX-Tol2* (Almeida et al., 2011b) at 25 ng/ul into single cell stage zebrafish embryos. Embryos screened at either 4 dpf or 5 dpf for EGFP-CAAX expression mounted in low melting termperature agarose and immersed in embryo medium containing Tricaine and imaged using a Zeiss Cell Observer SD 25 spinning disk confocal system spinning disk microscope equipped with a C-Apochromat 40X/1.1 NA water immersion objective. For *ntrk2a* mutant sheath analysis, petri dishes of embryos were blinded before imaging, followed by genotyping. For *bdnf* mutant sheath analysis, images were blinded using a lab utility file blinding plug in on ImageJ after genotyping. Prior to sheath measurements, images were deconvoluted using Zen 2.6 Blue Edition software using the regularized inverse filter (better, fast) deconvolution. Sheath measurements were obtained using the simple neurite tracer neuroanatomy ImageJ plugin, and we used Microsoft excel to compile cumulative sheath lengths (sum), average sheath lengths, and number of sheaths per cell for later analysis in Prism 9.9.

## Author Contributions

B.A. designed research, K.R., C.K., M.W., C.K., and C.A.D. performed research and analyzed data, A.R. and B.A. provided funding, and B.A. wrote the paper. All authors read and edited the manuscript.

The authors declare no conflict of interest.

## Acknowledgments

We thank members of the Appel lab for discussions and our zebrafish facility staff for excellent animal care. This work was supported by National Institutes of Health (NIH) grants R01NS065788, R01NS095679, and R35NS122191 to B.A., and R01NS086839 to A.R., and a gift from the Gates Frontiers Fund to B.A.

